# Haplotype Diversity and Sequence Heterogeneity of Human Telomeres

**DOI:** 10.1101/2020.01.31.929307

**Authors:** Kirill Grigorev, Jonathan Foox, Daniela Bezdan, Daniel Butler, Jared J. Luxton, Jake Reed, Cem Meydan, Susan M. Bailey, Christopher E. Mason

**Affiliations:** Department of Physiology and Biophysics, Weill Cornell Medicine, New York, New York, USA; The HRH Prince Alwaleed Bin Talal Bin Abdulaziz Alsaud Institute for Computational Biomedicine, Weill Cornell Medicine, New York, New York, USA; The Feil Family Brain and Mind Research Institute, New York, New York, USA; Department of Environmental and Radiological Health Sciences, Colorado State University, Fort Collins, CO; Cell and Molecular Biology Program, Colorado State University, Fort Collins, CO; The WorldQuant Initiative for Quantitative Prediction, Weill Cornell Medicine, New York, NY, USA

## Abstract

Telomeres are regions of repetitive nucleotide sequences capping the ends of eukaryotic chromosomes that protect against deterioration, whose lengths can be correlated with age and disease risk factors. Given their length and repetitive nature, telomeric regions are not easily reconstructed from short read sequencing, making telomere sequence resolution a very costly and generally intractable problem. Recently, long-read sequencing, with read lengths measuring in hundreds of Kbp, has made it possible to routinely read into telomeric regions and inspect their structure. Here, we describe a framework for extracting telomeric reads from single-molecule sequencing experiments, describing their sequence variation and motifs, and for haplotype inference. We find that long telomeric stretches can be accurately captured with long-read sequencing, observe extensive sequence heterogeneity of human telomeres, discover and localize non-canonical motifs (both previously reported as well as novel), and report the first motif composition maps of human telomeric diplotypes on a multi-Kbp scale.

## Introduction

Telomeres are the functional ends of human chromosomes that naturally shorten with mitosis and age [1], whose lengths can also be influenced by disease and environmental exposures (e.g., radiation, pollution, exercise, cancers) [2]. While human telomeres are known to consist largely of a conserved six-nucleotide repeat (TTAGGG) [3], several studies have identified variations of this motif in proximal telomeric regions [4–7]. However, such studies were performed with oligonucleotide hybridization, PCR, immunoprecipitation, and short read sequencing, resulting in discovery, but not localization, of motif variants. Thus, long-range maps of telomeric sequence variation in the human genome are still lacking. Such maps can provide insight into telomere biology and enable novel approaches to analyze the effects of aging, disease, and environment on telomere structure and length.

To improve our understanding of telomere structure and sequence variation, we developed *edgeCase*, a framework for alignment, motif discovery, and haplotype inference from human telomeric reads. We have validated these methods using Genome in a Bottle [8] single-molecule real-time (SMRT) sequencing datasets generated with Pacific Biosciences circular consensus sequencing (PacBio CCS) [9, 10] and short read Illumina [11] datasets. These results provide evidence for multiple novel, non-canonical telomeric repeats, resolution of chromosome-specific diplotypes with SMRT sequencing, and a new method for long-range characterization of the structure of telomeric sequences.

## Results

### Telomeric reads are present in human long-read whole genome sequencing datasets

We aligned PacBio CCS reads of three Genome in a Bottle (GIAB) human subjects (HG001, HG002, and HG005) to a combination of the human reference genome and human subtelomeric assemblies (see Materials and Methods). In total, we observed reads mapping to the ends of chromosomes and extending past them into telomeric regions on 9 *p* arms and 17 *q* arms, with 256 such reads (~10x mean coverage) in the HG001 dataset, 570 (~22x) in HG002, and 241 (~9x) in HG005. Figure 1 schematically represents the alignment of such reads in the HG002 dataset;alignment plots for the other two datasets are available as a supplementary figure (Figure S1), and full mapping statistics are available in Table S1. Illumina reads from matching GIAB datasets supported 70.8%, 63.3%, and 82.7% of the candidate PacBio CCS sequence, providing average coverages of ~5x, ~9x, and ~6x, respectively, by sequences supported by both technologies.

**Figure 1:**
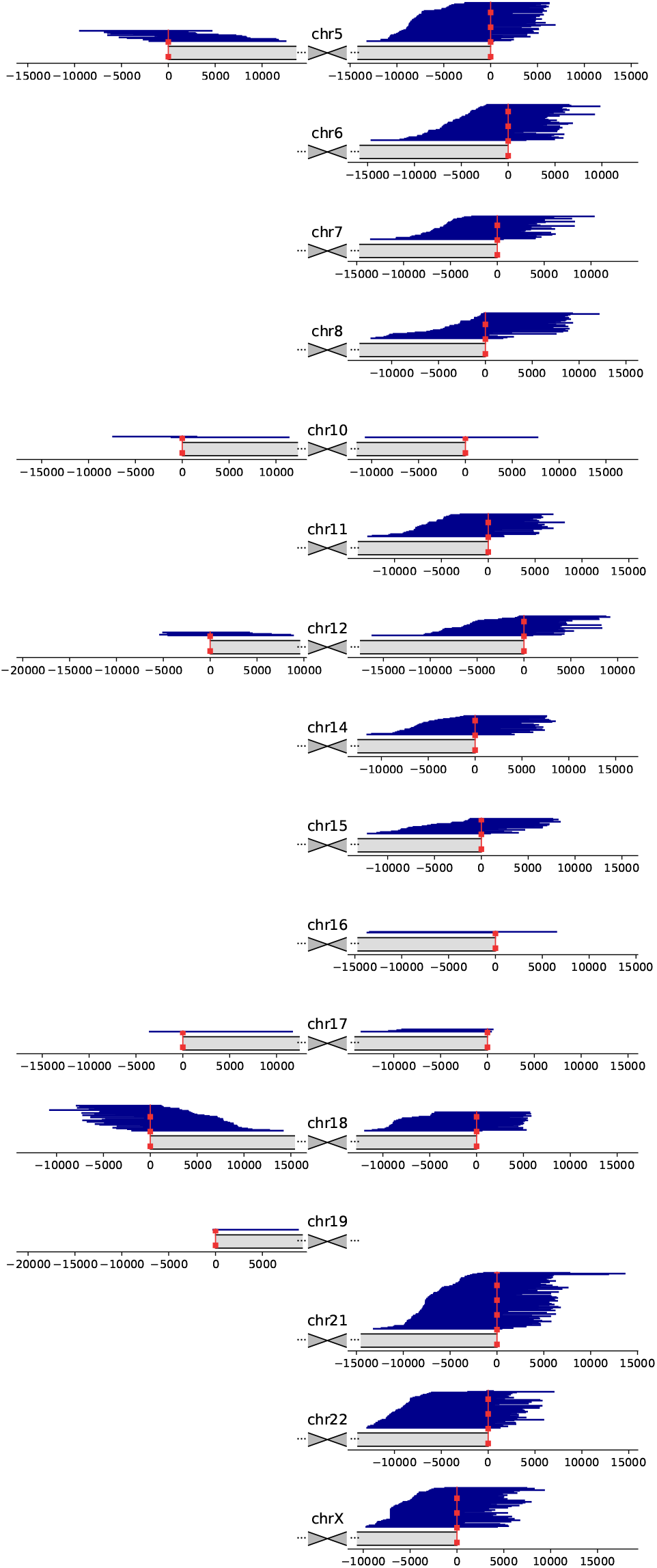
Mapping of candidate telomeric PacBio CCS reads from the HG002 dataset. Chromosomes are displayed schematically, centered around the centromere, with only the arms shown to which candidate reads aligned. Vertical red dashed lines denote the position of the boundary of the annotated telomeric tract. Coordinates are given in bp, relative to the positions of the telomeric tract boundaries.

### Telomeric reads contain variations of the canonical motif

We performed *de novo* repeat discovery in the supported regions for motifs of lengths 4 through 16 and identified motifs in repeat contexts that are statistically enriched in all three datasets. The majority of motifs were either the canonical TTAGGG/CCCTAA, its variation (e.g., TT<u>G</u>GGG/CCC<u>C</u>AA), or a duplet of variants, such as TTAGGGTTA<u>G</u>GGG (Table 1). CG-rich motifs were also observed on the *p* arms. When the entire candidate PacBio CCS sequences were considered, the top enriched motif (TTAGGG) explained 79.9% of the telomeric component, while all enriched motifs combined explained 87.7%. Figure 2 visualizes the locations of top four enriched motifs on the *q* arm of the HG002 dataset;only the arms covered by at least 20 reads are displayed. Plots for other datasets and arms are available as supplementary figures: Figure S2 visualizes the motifs on the *p* arm of the HG002 dataset, Figure S3 and Figure S4 visualize datasets HG001 and HG005 respectively. Reads on each arm agreed on the locations of different motifs within any given 10 bp window (the median of normalized Shannon entropy was 0.000 for all data, and the 3rd quartile was 0.166,0.074, and 0.211 for the three datasets, respectively, Figure S5), indicating that locations of the variations are colinear among reads and are not a result of sequencing errors.

**Figure 2:**
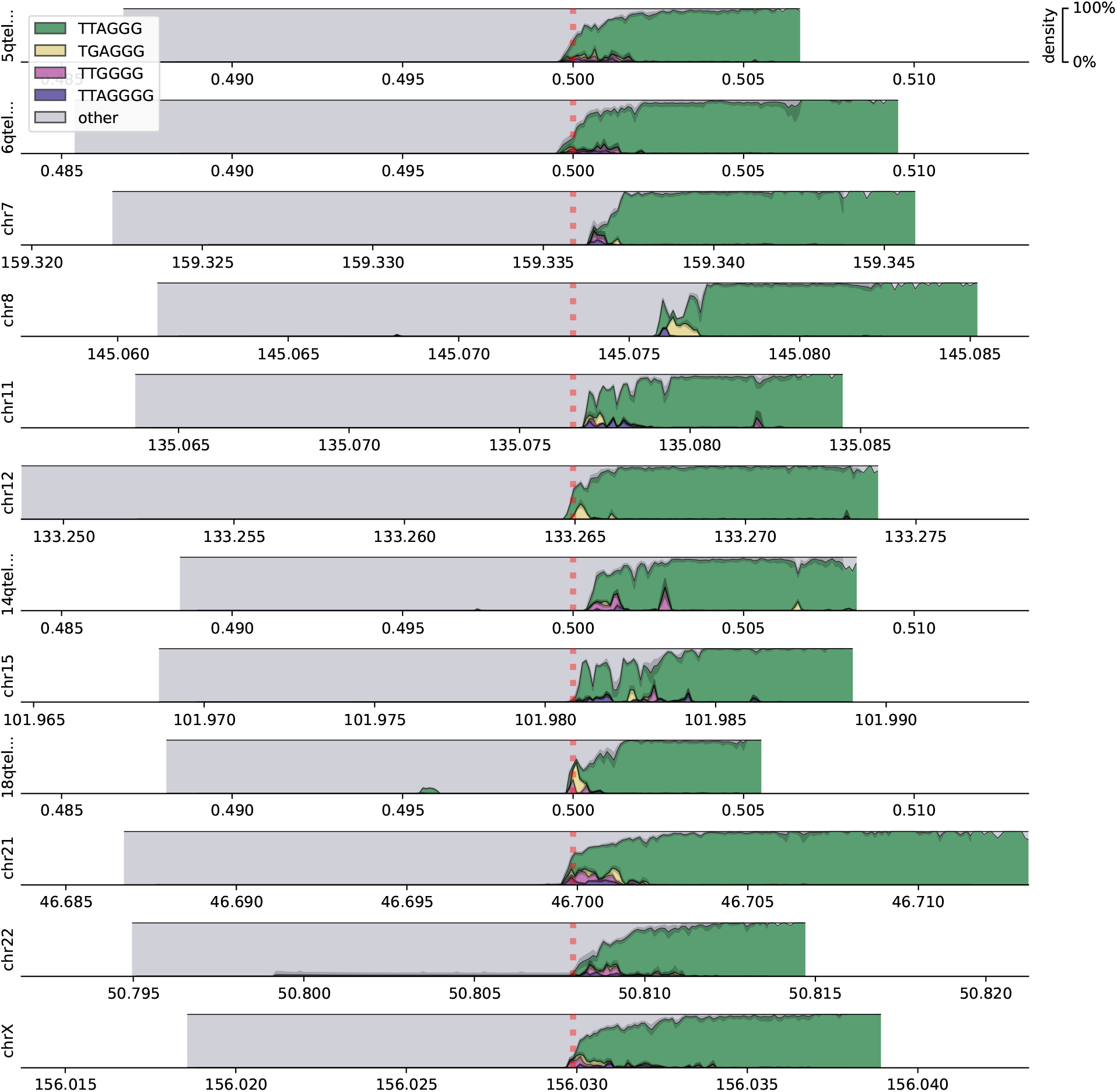
Motif densities at ends of chromosomal *q* arms of the HG002 dataset. Only the arms covered by at least 20 reads are displayed. Shaded boxes span the mapped regions of the genome. Motif densities are plotted as stacked area charts;ribbons surrounding area boundaries represent the 95% confidence interval of bootstrap. Top four enriched motifs are plotted in color;pale tinted areas represent the density of any other motifs and non-repeating sequences (absence of enriched motifs). Absolute genomic coordinates are given in Mbp on the specific reference contigs the reads mapped to (for example, for chr5, reads mapped to the 500 Kbp-long subtelomeric assembly 5qtel_1-500K_1_12_12). Vertical red dashed lines denote the position of the boundary of the annotated telomeric tract.

**Table 1:**
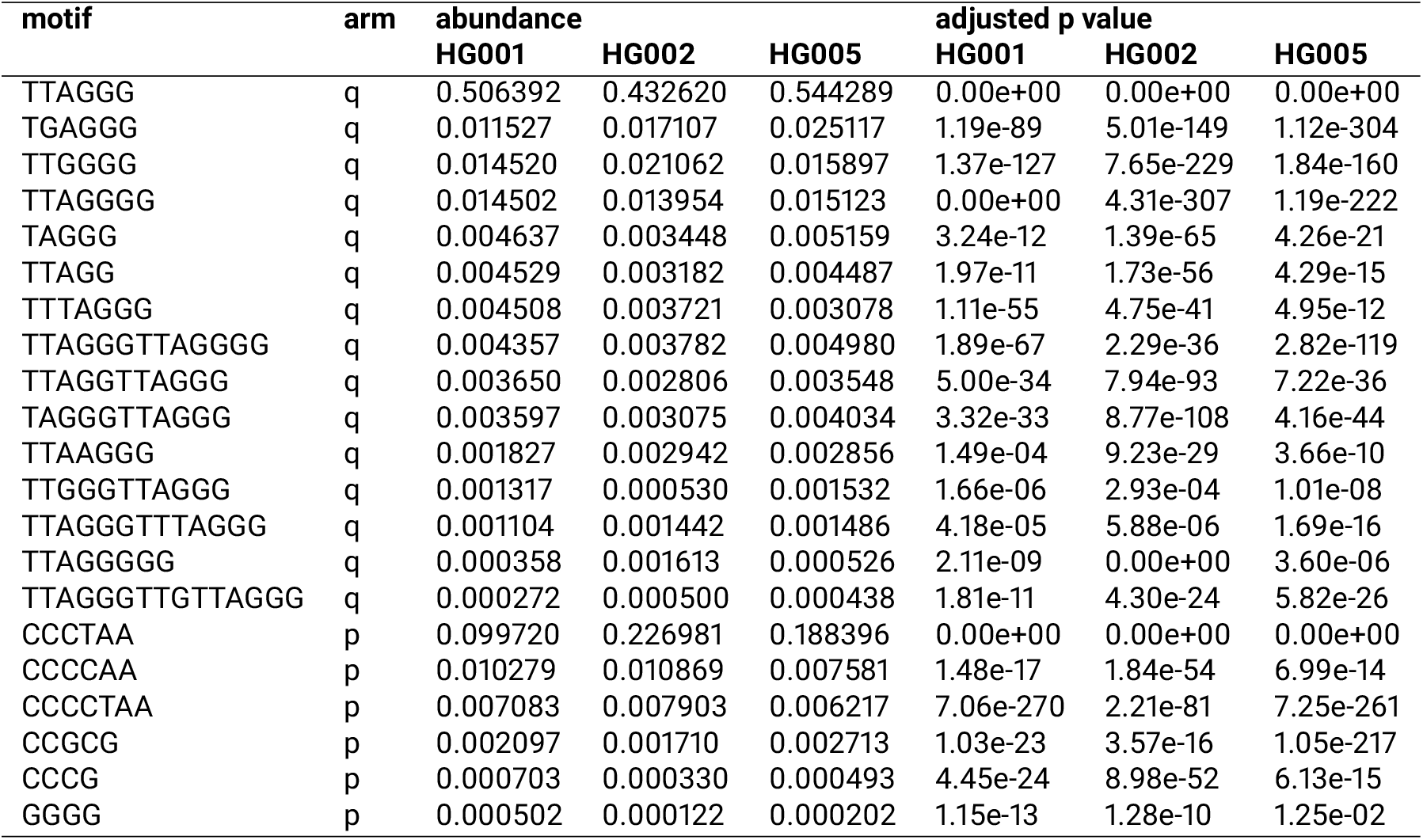
Significantly enriched repeating motifs in telomeric regions of GIAB datasets HG001, HG002, and HG005.

### Long-read sequencing resolves human telomeric haplotypes

Sequences of telomeric reads clustered by relative pairwise Levenshtein distances [12] with varying levels of heterogeneity depending on the dataset and the chromosomal arm to which they belonged. We examined the *q* arms of the HG002 dataset to investigate this heterogeneity, as they provided the deepest coverage (Table S1), and found that, on 12 out of the 15 arms, reads clustered into two prominent groups per arm when maximizing the Bayesian information criterion [13] (see Materials and Methods). Pairwise distances between the reads within these clusters were significantly lower than those for out-of-cluster pairings, implying that distinct telomeric haplotypes are present. To quantify the differences between putative haplotypes, we calculated silhouette scores [14] for these clusterings (Table 2), and generated motif density plots for the four chromosome arms with the highest such scores to visualize the differences in haplotypes (Figure 3).

**Figure 3:**
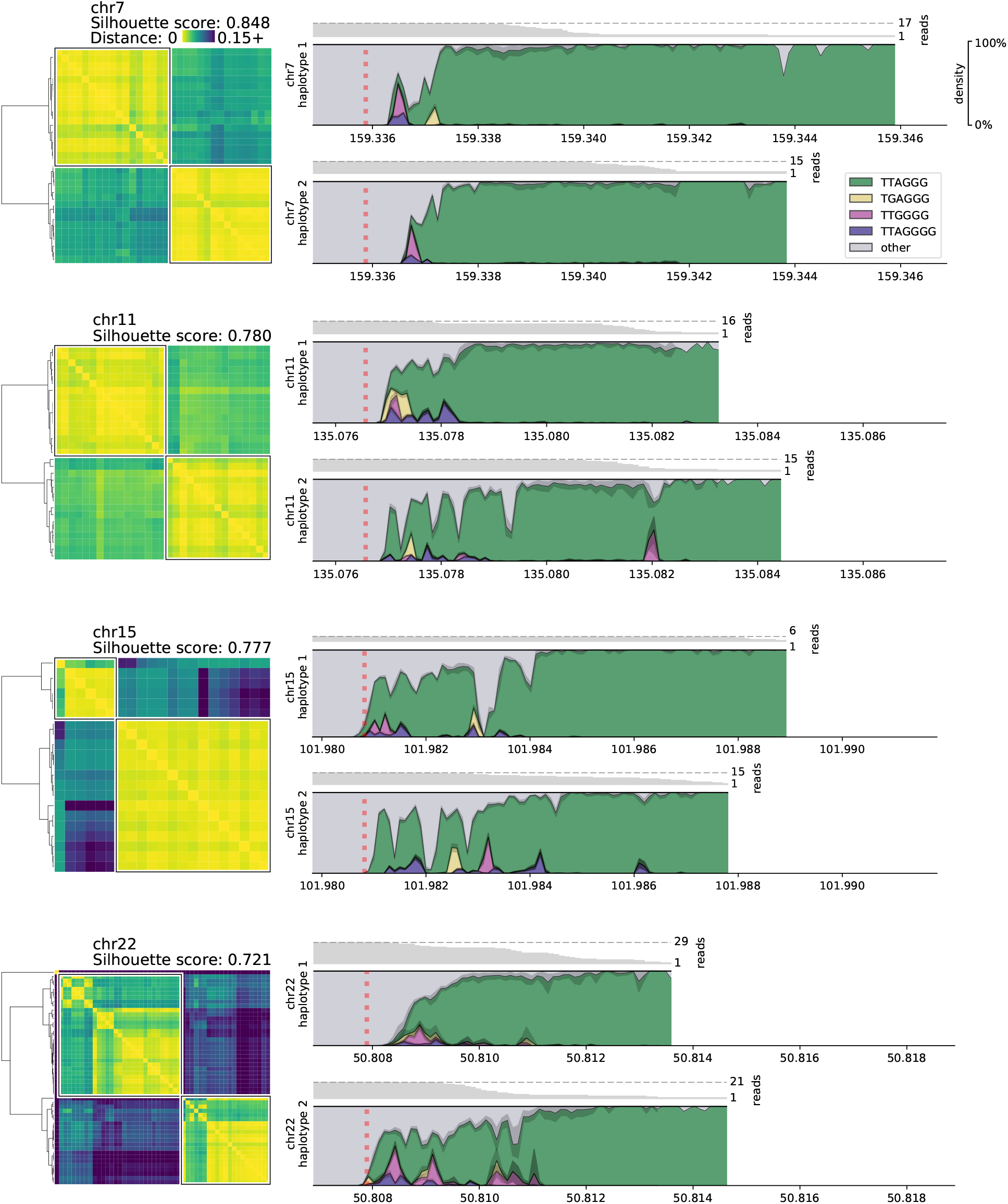
Clustering of reads into haplotypes based on relative pairwise Levenshtein distances on four representative chromosomal *q* arms in the HG002 dataset, and densities of top enriched motifs in each haplotype. Genomic coordinates are given in Mbp. Read coverage of each haplotype is annotated above the density plot.

**Table 2:**
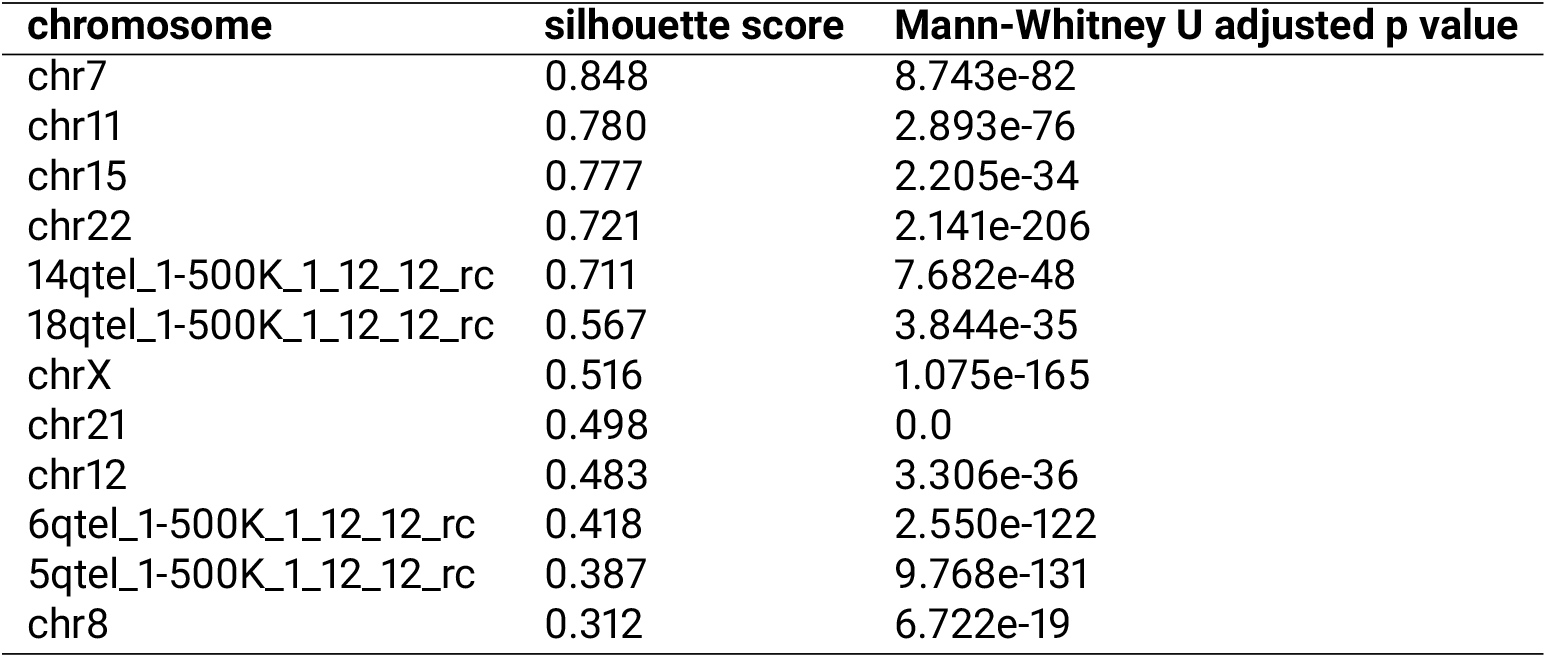
Silhouette scores for cluster assignments on *q* arms of chromosomes in sample HG002 where two clusters of reads are present, and adjusted *p*-values for the Mann-Whitney U test for the difference of within-cluster and out-of-cluster relative pairwise Levenshtein distances.

## Discussion

Repeat-rich, low-complexity regions of the human genome such as telomeres have been historically recalcitrant to full mapping and annotation [15], mainly due to the alignment challenge they pose and to the read lengths required to span such areas [16]. The advent of long-read, single-molecule methods (third generation sequencing) has provided new opportunities to map the sequence composition of a previously “dark” area of the human genome. These results reaffirm that the canonical repeat (TTAGGG) is certainly the most dominant type of motif in telomeres, but also reveal a surprising diversity of repeat variations, which are confirmed by both short and long-read sequencing technologies. This diversity of repeats includes previously reported variants, as well as novel motifs that are characterized not only by nucleotide substitutions, but also insertions, deletions, and even motif pairing. Apart from these variations, CG-rich motifs were identified in telomeric regions of *p* arms, consistent with previously reported findings [17];interestingly, CG content was more pronounced in subtelomeric regions immediately adjacent to the telomere, which may have implications for epigenetic regulation. Moreover, while short read sequencing is able to identify such variants, it alone cannot reveal the relative locations of these motifs within telomeres, as repetitive short reads can neither be aligned outside of the reference genome nor provide enough overlap variability to be assembled *de novo.* Long SMRT reads, on the other hand, can be anchored to known subtelomeric sequences of the human genome and extend into the previously unmapped telomeric area. These results also highlight the need of better subtelomeric and telomeric annotations in the human genome. Four of the 40 subtelomeric assemblies [18] were homologous to regions in the reference genome far within the respective chromosomes (up to 586 Kbp into the reference sequence), and the canonical motif was present on the *q* arm of chr8 only after 2-3Kbp past the annotated boundary in all datasets, suggesting that the existing assemblies do not provide a completely accurate telomeric annotation, and that methods described herein could help to resolve these areas of reference genomes.

We observed PacBio CCS reads reaching up to 13 Kbp beyond the known regions of the genome, and resolving the underlying sequence with reasonable fidelity - even without support from short reads, - both measured by the entropy of motif assignment and by pairwise Levenshtein distances between the reads belonging to the same chromosomal arms. While Illumina reads also provided support for all of the reported motifs, the overlap between the short and the long reads was substantial, but not complete, which can be explained by the necessary bias towards the canonical motif during the selection of short reads. Therefore, telomeric regions with higher content of non-canonical repeats are less likely to be identified through the use of short reads, and instead, long reads appear to be more suitable for this purpose as well. The identified variations in long range contexts enable clustering of SMRT reads into distinct haplotypes at ends of chromosomes, and thus provide a new means of diplotype mapping and reveal the existence and motif composition of such diplotypes on a multi-Kbp scale.

## Materials and Methods

### The extended reference genome

We constructed the extended reference genome by performing an all-to-all alignment of all contigs in the *hg38* reference genome [19,20] and the subtelomeric assemblies [18] with *minimap2* [21] using three settings for assembly-to-reference mapping (*asm5, asm10, asm20*). Forty subtelomeric contigs mapped to ends of *hg38* chromosomes with a mapping quality of 60, one (XpYptel) mapped with the quality of 0 and was discarded; one (14qtel) mapped to the ALT version of chr14 (chr14_KI270846v1_alt) with the quality of 52, which, in turn, mapped to the main chr14 chromosome with the quality of 60. These data and the exact match and mismatch coordinates were used to create a combined reference *(hg38ext)* in which subtelomeric contigs informed the locations of the boundaries of the telomeric tracts *(tract_anchor).* Such contigs that mapped fully within *hg38* chromosomes resulted in *tract_anchor* annotations directly on those *hg38* chromosomes; partially mapping contigs were considered as forking from the *hg38* sequence and were similarly annotated by themselves.

### Selection of telomeric reads and identification of repeat content

Three subjects were selected for the analysis. The first individual (NA12878IHG001) came from the pilot genome of the HapMap project [22], while the other two, including the son from the Ashkenazi Jewish Trio (NA24385IHG002) and the son from the Chinese Trio (NA24631IHG005), are members of the Personal Genome Project, whose genomes are consented for commercial redistribution and reidentification [23]. These subjects are referred to hereafter as HG001, HG002, and HG005, respectively.

For subjects HG001 and HG005, Genome in a Bottle [8] PacBio_SequelII_CCS_11kb datasets were used (one dataset per each subject). For subject HG002, a combination of two sequencing experiments was analyzed (PacBio_CCS_10kb and PacBio_CCS_15kb). The mean coverage was ~29x, ~58x, and ~32x for subjects HG001, HG002, and HG005, respectively. Reads were mapped to *hg38ext* with *minimap2*, and reads that mapped to either end of either chromosome and overlapped the boundary of its telomeric tract were selected for further analysis. These reads had a portion of their sequence mapped to the reference contig and a portion extending beyond the reference (soft- or hard-clipped in the alignment file). Sequences past the *tract_anchor* marker were extracted from the reads that had this marker within their mapped portion (from the 5’ end to the marker on *p* arms and from the marker to the 3’ end on *q* arms, accounting for forward and reverse mappings). To identify regions of the telomeres that are fully supported by both short and long reads, we extracted candidate telomeric reads from GIAB Illumina datasets (NIST_NA12878_-HG001_HiSeq_300x, NIST_HiSeq_HG002_Homogeneity-10953946, HG005_NA24631_son_HiSeq_300x;all three ~300x coverage) with *Telomerecat* [24], and selected those that mapped perfectly with *minimap2* (at least a 50bp-long exact match without insertions or deletions, allowing all secondary mappings) to the telomeric regions of the PacBio CCS candidates from the same subject’s dataset.

Within the regions supported by both PacBio CCS and Illumina candidate reads, overrepresentation of motifs of lengths *k* ⊂ [4..16] was tested. To target motifs in repeat contexts and to distinguish motifs differing by indels (for example, ACGT and ACGTT), doubled sequences (for example, *k*-mer ACGTACGT for motif ACGT) were counted with *jellyfish* [25], and counts of *k*-mers synonymous with respect to circular shifts (for example, ACGTACGT and CGTACGTA) were summed together. For each such *k*-mer, Fisher’s exact test was performed to determine whether its count is significant on the background of counts of other *k*-mers of the same length. Briefly, we considered *k*-mers with counts higher than 1.5 interquartile range above the third quartile of the distribution as potentially classifiable, and a 2×2 contingency matrix *C* for the test was constructed as follows: row 0 contained counts of potentially classifiable *k*-mers, row 1 contained counts of remaining (non-classifiable) *k*-mers, columns 0 and 1 contained counts of single and remaining (background) *k*-mers, respectively, i.e.: *C*_0,0_ = count of target *k*-mer, *C*_0,1_ = sum of counts of other potentially classifiable *k*-mers, *C*_1,0_ = median count of *k*-mer, *C*_1,1_ = sum of counts of other non-classifiable *k*-mers. The Bonferroni multiple testing correction was applied to the resultant *p*-values, and motifs for which *k*-mers yielded *p*-values below the cutoff of 0.05 were reported.

As telomeric reads contain long low-complexity regions and present an alignment challenge, we evaluated concordance of their sequences without realignment of their portions that extended past the reference sequence. To that end, for all reads mapping to the same chromosomal arm, we calculated densities of each motif in a rolling window starting from the innermost mapped position. To evaluate whether the reads on the same arm agree on the positions of different motifs, for each read, we calculated motif densities in 10 bp windows with 10 bp smoothing to buffer insertions and deletions. For each window in a read, the motif with the highest density was selected to represent that window. Then, normalized Shannon entropy among all reads was calculated in each window as 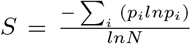, where *p_i_* is the frequency of each motif in the window and *N* is the number of motifs [26]. The value of normalized entropy was a metric bounded by [0,1], with 0 describing perfect agreement and 1 describing maximum randomness. For visualization, we performed 1000 rounds of bootstrap of the calculated density values in 100 bp rolling windows, and selected the lower and the upper bounds of the 95% confidence interval of bootstrap. Of note, several chromosome arms had the *tract_anchor* position further away from the end of the contig than others (~79-586 Kbp into the chromosome sequence), and the reads mapping to these arms did not contain these motifs, suggesting that either their subtelomeric annotations were incorrect or large insertions or duplications were present in the reference genome;in light of this, reads mapping to the *p* arm of chr1, the *q* arm of chr4, and both arms of chr20 were removed from the study, and the analysis was repeated.

### Extraction of telomeric haplotypes

Within groups of reads mapping to each chromosome arm, all relative pairwise Levenshtein distances were calculated. In short, to calculate the absolute distance between each pair of reads, the sequences in the overlapping positions of the reads were extracted;the distance then equaled the minimum number of single-character insertions, deletions, and substitutions required to make these sequences identical. The relative distance was computed as the absolute distance divided by the length of the overlap. Relative distances were then clustered using Ward’s method via the Euclidean metric. The optimal number of clusters was determined by maximizing the Bayesian information criterion [13], allowing for no more than one outlier and at least five reads per cluster, and silhouette scores for these clusterings were calculated. Briefly, as previously described [14], a silhouette score of a clustering was computed as the mean value of silhouette coefficients of all entries, which, in turn, equaled 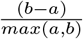 where *a* = mean intra-cluster distance and *b* = mean nearest-cluster distance for an entry. Levenshtein distances of all within-cluster pairings and of all out-of-cluster pairings were compared using the one-tailed Mann-Whitney U test; *p*-values were adjusted with the Bonferroni correction. Distinct clusters of reads within the same chromosome arm (adjusted Mann-Whitney U *p*-value below 0.05) were reported as putative haplotypes. As the HG002 dataset was combined from two sequencing experiments, we investigated the provenance of reads in these haplotypes;reads from both sequencing experiments contributed to each haplotype with an average ~1:2 ratio (Table S2).

## Availability and implementation

The software for identification of telomeric reads, *de novo* discovery of repeat motifs, haplotype inference and motif density visualization was implemented in Python and is freely available at github.com/lankycyril/edgecase.

## Acknowledgements

We would like to thank the Epigenomics Core Facility at Weill Cornell Medicine, the Scientific Computing Unit (SCU), XSEDE Supercomputing Resources, as well as the STARR grants I9-A9-071, I13-0052, The Vallee Foundation, The WorldQuant Foundation, The Pershing Square Sohn Cancer Research Alliance, NASA (NNX14AH51G, NNX14AB02G, NNX17AB26G), The National Institutes of Health (R01MH117406, R01NS076465, R01CA249054, R01AI151059, P01HD067244, P01CA214274), TRISH (NNX16AO69A:0107, NNX16AO69A:0061), the LLS (9238-16, Mak, MCL-982, Chen-Kiang), and the NSF (1840275).

## Author contributions

S.M.B. and C.E.M. conceived the study. K.G., J.F., and C.E.M. developed the framework and analyzed the data. D.Be., D.Bu., J.J.L, J.R., and C.M. analyzed the data. All authors edited the manuscript.

## Supplementary materials

### Supplementary figures

**Figure S1:**
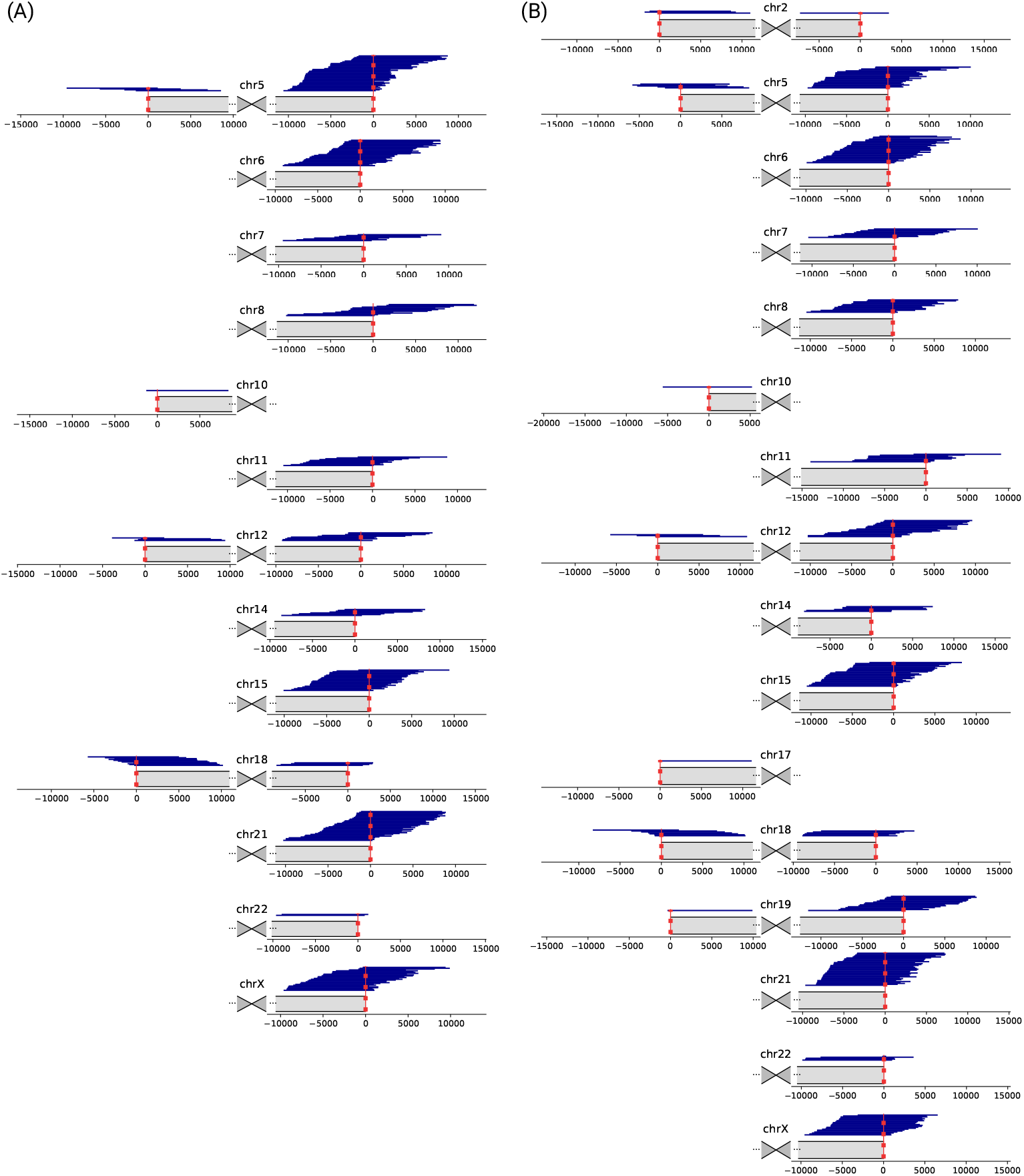
Mapping of candidate telomeric PacBio CCS reads from datasets (A) HG001 and (B) HG005. Chromosomes are displayed schematically, centered around the centromere, with only the arms shown to which candidate reads aligned. Vertical red dashed lines denote the position of the boundary of the annotated telomeric tract. Coordinates are given in bp, relative to the positions of the telomeric tract boundaries.

**Figure S2:**
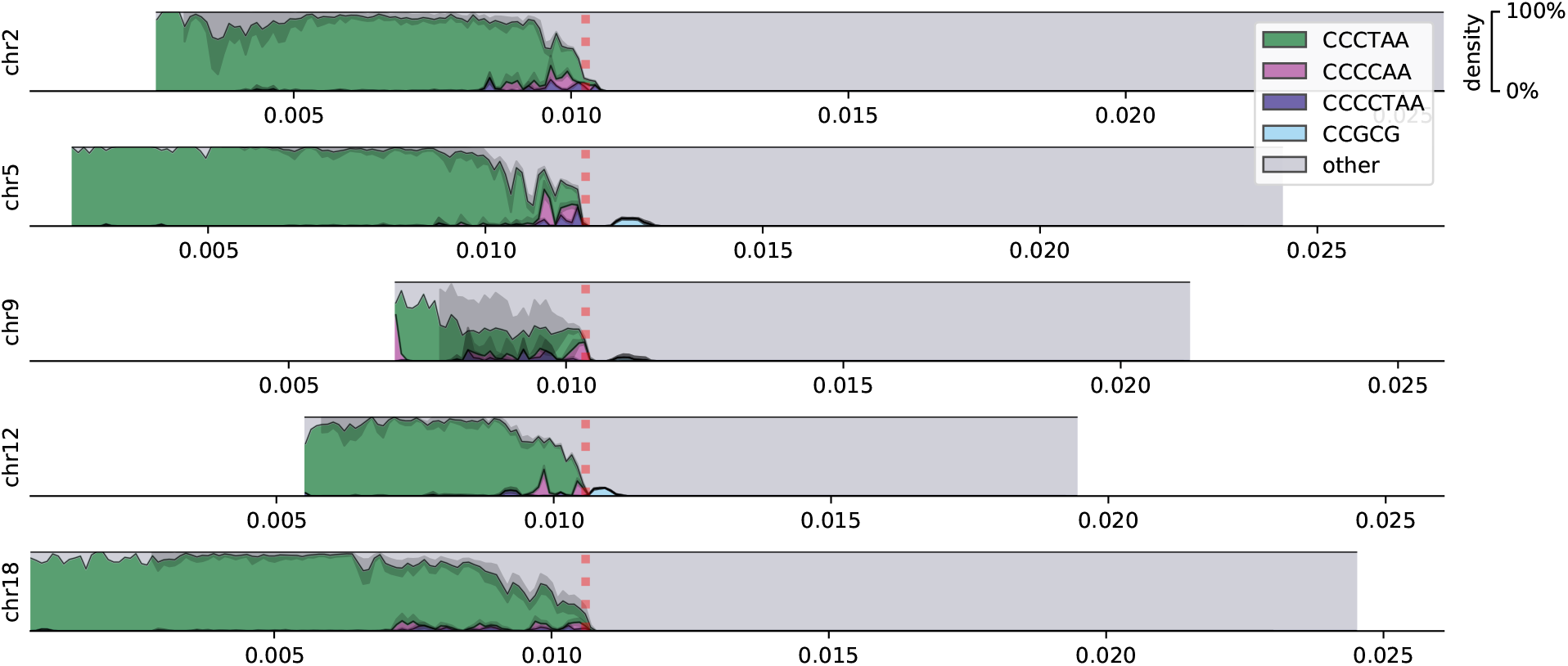
Motif densities at ends of chromosomal *p* arms of the HG002 dataset. Only the arms covered by at least 20 reads are displayed. Genomic coordinates are given in Mbp. Vertical red dashed lines denote the position of the boundary of the annotated telomeric tract. The density of the CCGCG motif falls below the resolution of visual detection in the telomeric tract, but is also present in the subtelomeric region.

**Figure S3:**
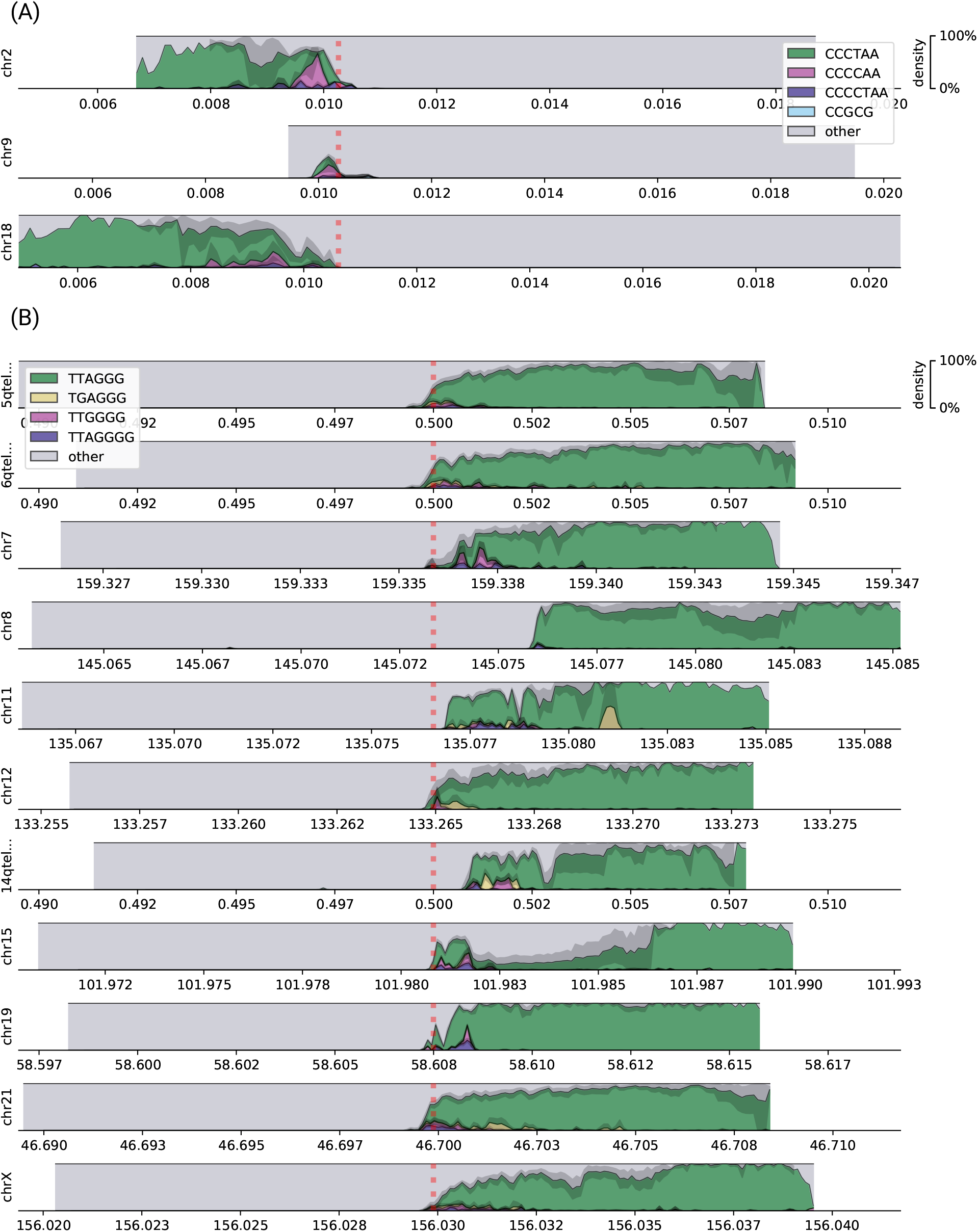
Motif densities at ends of chromosomal (A) *p* and (B) *q* arms of the HG001 dataset. Only the arms covered by at least 20 reads are displayed. Genomic coordinates are given in Mbp.

**Figure S4:**
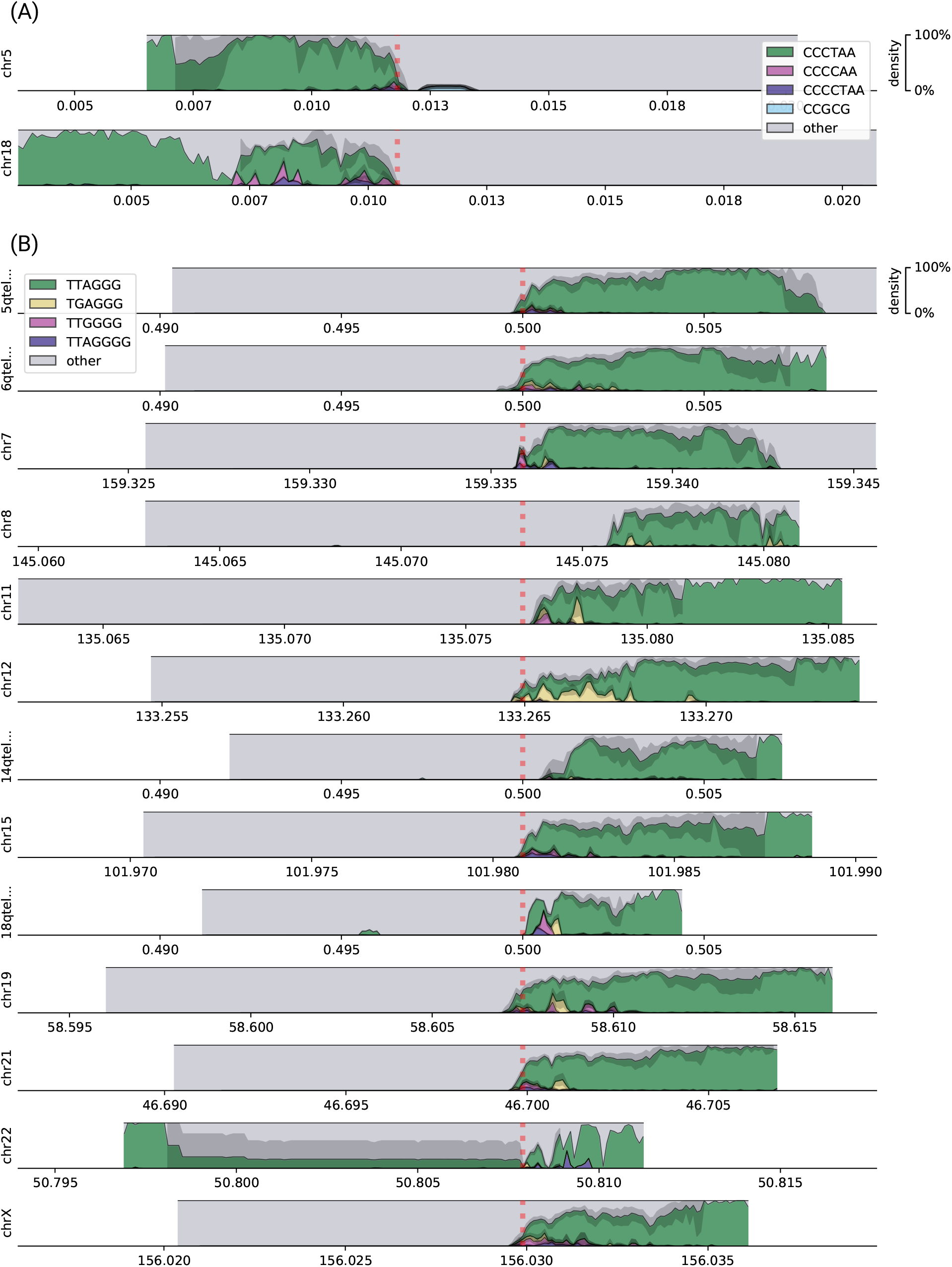
Motif densities at ends of chromosomal (A) *p* and (B) *q* arms of the HG005 dataset. Only the arms covered by at least 20 reads are displayed. Genomic coordinates are given in Mbp.

**Figure S5:**
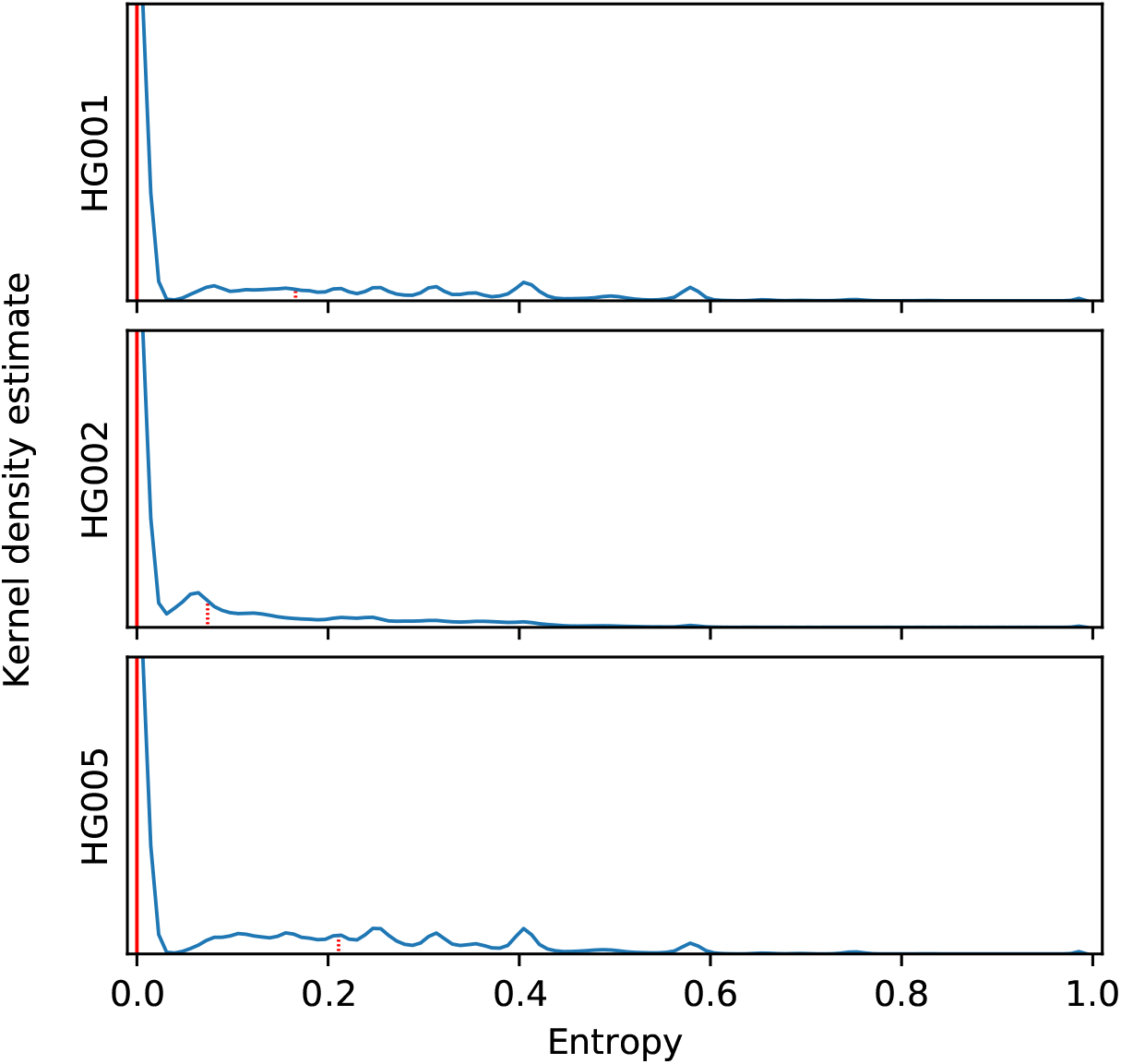
Distribution of motif entropies in 10 bp windows of candidate PacBio CCS reads aligning to the same chromosomal arms in GIAB datasets HG001, HG002, and HG005. Red solid lines denote the position of the median (0.000 in all three datasets), and red dashed lines denote the 3rd quartile (0.166, 0.074, and 0.211, respectively).

### Supplementary tables

**Table S1:** The number of telomeric reads on each arm identified in GIAB PacBio CCS datasets HG001, HG002, and HG005.

| chromosome | reference contig | arm | HG001 | HG002 | HG005 |
| --- | --- | --- | --- | --- | --- |
| chr2 | 2qtel_1-500K_1_12_12_rc | q | 0 | 0 | 1 |
| chr2 | chr2 | p | 5 | 16 | 3 |
| chr5 | 5qtel_1-500K_1_12_12_rc | q | 42 | 53 | 23 |
| chr5 | chr5 | p | 4 | 15 | 5 |
| chr6 | 6qtel_1-500K_1_12_12_rc | q | 31 | 49 | 29 |
| chr7 | chr7 | q | 8 | 32 | 10 |
| chr8 | chr8 | q | 14 | 35 | 14 |
| chr9 | chr9 | p | 6 | 6 | 0 |
| chr10 | 10qtel_1-500K_1_12_12_rc | q | 0 | 1 | 0 |
| chr10 | chr10 | p | 1 | 2 | 1 |
| chr11 | chr11 | q | 11 | 31 | 9 |
| chr12 | chr12 | q | 10 | 27 | 18 |
| chr12 | chr12 | p | 4 | 5 | 3 |
| chr14 | 14qtel_1-500K_1_12_12_rc | q | 8 | 26 | 6 |
| chr15 | chr15 | q | 25 | 21 | 26 |
| chr16 | 16qtel_1-500K_1_12_12_rc | q | 0 | 2 | 0 |
| chr16 | chr16 | p | 1 | 0 | 0 |
| chr17 | 17qtel_1-500K_1_12_12v2_rc | q | 0 | 4 | 0 |
| chr17 | 17ptel_1_500K_1_12_12 | p | 0 | 1 | 1 |
| chr18 | 18qtel_1-500K_1_12_12_rc | q | 4 | 26 | 6 |
| chr18 | chr18 | p | 11 | 35 | 7 |
| chr19 | 19ptel_1-500K_1_12_12 | p | 0 | 1 | 1 |
| chr19 | chr19 | q | 6 | 0 | 16 |
| chr21 | chr21 | q | 35 | 77 | 35 |
| chr22 | chr22 | q | 2 | 51 | 5 |
| chrX | chrX | q | 28 | 54 | 22 |

**Table S2:** Amounts of reads from the two HG002 PacBio CCS sequencing experiments contributing to each telomeric haplotype on the *q* arms.

| chromosome | haplotype | PacBio_CCS_10kb | PacBio_CCS_15kb |
| --- | --- | --- | --- |
| 5qtel_1-500K_1_12_12_rc | 1 | 11 | 24 |
| 5qtel_1-500K_1_12_12_rc | 2 | 7 | 10 |
| 6qtel_1-500K_1_12_12_rc | 1 | 8 | 12 |
| 6qtel_1-500K_1_12_12_rc | 2 | 18 | 10 |
| chr7 | 1 | 9 | 8 |
| chr7 | 2 | 7 | 8 |
| chr8 | 1 | 8 | 6 |
| chr8 | 2 | 9 | 8 |
| chr11 | 1 | 5 | 11 |
| chr11 | 2 | 8 | 7 |
| chr12 | 1 | 9 | 9 |
| chr12 | 2 | 6 | 3 |
| 14qtel_1-500K_1_12_12_rc | 1 | 3 | 8 |
| 14qtel_1-500K_1_12_12_rc | 2 | 5 | 10 |
| chr15 | 1 | 6 | 0 |
| chr15 | 2 | 4 | 11 |
| 18qtel_1-500K_1_12_12_rc | 1 | 4 | 9 |
| 18qtel_1-500K_1_12_12_rc | 2 | 4 | 9 |
| chr21 | 1 | 16 | 20 |
| chr21 | 2 | 12 | 29 |
| chr22 | 1 | 2 | 27 |
| chr22 | 2 | 11 | 10 |
| chrX | 1 | 12 | 13 |
| chrX | 2 | 10 | 19 |

## References

1. Aubert, G. & Lansdorp, P. M. Telomeres and Aging. Physiological Reviews 88 (Apr. 2008).

2. Shammas, M. A. Telomeres, lifestyle, cancer, and aging. Current Opinion in Clinical Nutrition and Metabolic Care 14 (Jan. 2011).

3. Moyzis, R. K. et al. A highly conserved repetitive DNA sequence, (TTAGGG)n, present at the telomeres of human chromosomes. Proceedings of the National Academy of Sciences 85 (Sept. 1988).

4. Allshire, R. C., Dempster, M. & Hastie, N. D. Human telomeres contain at least three types of G-rich repeat distributed non-randomly. Nucleic Acids Research 17 (1989).

5. Coleman, J., Baird, D. M. & Royle, N. J. The Plasticity of Human Telomeres Demonstrated by a Hypervariable Telomere Repeat Array That Is Located on Some Copies of 16p and 16q. Human Molecular Genetics 8 (Sept. 1999).

6. Lee, M. et al. Telomere sequence content can be used to determine ALT activity in tumours. Nucleic Acids Research 46 (Apr. 2018).

7. Bluhm, A. et al. ZBTB10 binds the telomeric variant repeat TTGGGG and interacts with TRF2. Nucleic Acids Research 47 (Jan. 2019).

8. Zook, J. M. et al. An open resource for accurately benchmarking small variant and reference calls. Nature Biotechnology 37 (Apr. 2019).

9. Eid, J. et al. Real-Time DNA Sequencing from Single Polymerase Molecules. Science 323 (Jan. 2009).

10. Ardui, S., Ameur, A., Vermeesch, J. R. & Hestand, M. S. Single molecule real-time (SMRT) sequencing comes of age: applications and utilities for medical diagnostics. Nucleic Acids Research 46 (Feb. 2018).

11. Bentley, D. R. et al. Accurate whole human genome sequencing using reversible terminator chemistry. Nature 456 (Nov. 2008).

12. Levenshtein, V. I. Binary codes capable of correcting deletions, insertions, and reversals in Soviet physics doklady 10 (1966).

13. Schwarz, G. Estimating the Dimension of a Model. The Annals of Statistics 6 (Mar. 1978).

14. Finding Groups in Data (eds Kaufman, L. & Rousseeuw, P. J.) (John Wiley & Sons, Inc., Mar. 1990).

15. Miga, K. H. Completing the human genome: the progress and challenge of satellite DNA assembly. Chromosome Research 23 (Sept. 2015).

16. Treangen, T. J. & Salzberg, S. L. Repetitive DNA and next-generation sequencing: computational challenges and solutions. Nature Reviews Genetics 13 (Nov. 2011).

17. Nergadze, S. G. et al. CpG-island promoters drive transcription of human telomeres. RNA 15 (Oct. 2009).

18. Stong, N. et al. Subtelomeric CTCF and cohesin binding site organization using improved subtelomere assemblies and a novel annotation pipeline. Genome Research 24 (Mar. 2014).

19. Schneider, V. A. et al. Evaluation of GRCh38 and de novo haploid genome assemblies demonstrates the enduring quality of the reference assembly. Genome Research 27 (Apr. 2017).

20. Initial sequencing and analysis of the human genome. Nature 409 (Feb. 2001).

21. Li, H. Minimap2: pairwise alignment for nucleotide sequences. Bioinformatics 34 (May 2018).

22. The International HapMap Project. Nature 426 (Dec. 2003).

23. Zook, J. M. et al. Extensive sequencing of seven human genomes to characterize benchmark reference materials. Scientific Data 3 (June 2016).

24. Farmery, J. H. R., Smith, M. L. & Lynch, A. G. Telomerecat: A ploidy-agnostic method for estimating telomere length from whole genome sequencing data. Scientific Reports 8 (Jan. 2018).

25. Marçais, G. & Kingsford, C. A fast, lock-free approach for efficient parallel counting of occurrences of k-mers. Bioinformatics 27 (Jan. 2011).

26. Minosse, C. et al. Possible Compartmentalization of Hepatitis C Viral Replication in the Genital Tract of HIV-1-Coinfected Women. The Journal of Infectious Diseases 194 (Dec. 2006).

